# High temperature perception in leaves promotes vascular regeneration in distant tissues

**DOI:** 10.1101/2021.08.03.454483

**Authors:** Phanu T. Serivichyaswat, Kai Bartusch, Martina Leso, Constance Musseau, Akira Iwase, Yu Chen, Keiko Sugimoto, Marcel Quint, Charles W. Melnyk

**Affiliations:** Department of Plant Biology, Swedish University of Agricultural Sciences, Ulls gränd 1, 765 51 Uppsala, Sweden; Institute of Molecular Plant Biology, Department of Biology, ETH Zürich, Zurich, Switzerland; Institute of Agricultural and Nutritional Sciences, Martin Luther University Halle-Wittenberg, Betty-Heimann-Str. 5, 06120 Halle (Saale), Germany; RIKEN Center for Sustainable Resource Science, Yokohama 230-0045, Japan; Department of Biological Sciences, Faculty of Science, The University of Tokyo, Tokyo, 113-8654, Japan

**Author notes:** equal contribution (Phanu T. Serivichyaswat, Kai Bartusch). Correspondence: Charles W. Melnyk.

## Abstract

Cellular regeneration in response to wounding is fundamental to maintain tissue integrity. Various internal factors including hormones and developmental pathways affect wound healing but little is known about how external factors influence regeneration. To better understand how the environment affects regeneration, we investigated the effects of temperature using the horticulturally relevant process of plant grafting. We found that elevated temperatures accelerated vascular regeneration of *Arabidopsis thaliana* and tomato (*Solanum lycopersicum*) grafts. Leaves were critical for this effect since blocking auxin transport or mutating *PHYTOCHROME INTERACTING FACTOR4* (*PIF4*) or *YUCCA2/5/8/9* in the cotyledons abolished the temperature enhancement. However, these perturbations had no effect upon graft healing at ambient temperatures and mutations in *PIF4* did not affect the temperature enhancement of callus formation or tissue adhesion, suggesting that leaf-derived auxin was specific for enhancing vascular regeneration in response to elevated temperatures. Tissue-specific perturbations of auxin response using a *BODENLOS (BDL)* mutant revealed an asymmetric effect of temperature upon regeneration: the presence of *bdl* above the cut prevented temperature enhancement whereas the presence of *bdl* below the cut prevented graft healing regardless of temperature. Promotion of tissue regeneration by elevated temperatures was not specific for graft healing and we found that elevated temperatures accelerated xylem formation between the parasite *Phtheirospermum japonicum* and host *Arabidopsis thaliana*, and this effect required shoot-derived auxin from the parasite. Taken together, our results identify a pathway by which elevated temperatures accelerate vascular development which could be of relevance for improving regeneration and better understanding inter-plant vascular connections.

## Introduction

Various abiotic and biotic stresses including temperature extremes, herbivory and cutting induce cellular and tissue damage that needs to be repaired^1^. These environmental stresses are a source of tissue damage, but the environment can also promote regeneration. One notable example is the influence of temperature upon regeneration. Elevated temperatures enhance the formation of stem-cell like tissues termed callus that aid the wound healing process^2^. Increased temperatures also improve the horticultural process of plant grafting^3,4^ which consists of cutting and joining different plants together to obtain chimeric plants that improve stress tolerance and yields^5,6^. At the cut sites, grafts initially form callus^7^ that seal that wound, followed by vascular division and differentiation that allows phloem and xylem reconnection^8^. Similar processes occur during other forms of wound healing such as when callus forms at the site of cutting or underlying cell layers divide and differentiate to restore tissue integrity after cell ablation^9,10^. A common theme to regeneration in plants is the involvement of auxin. Auxin is mainly produced in young leaves^11^ and accumulates at the site of injury^12^ where auxin responses increase at both graft junctions and at the site of stem cutting or cell ablation^8,13,14^. Auxin plays an important role in regenerating the vasculature since disrupting auxin response below the graft junction inhibits graft formation^8^, whereas blocking auxin transport inhibits graft formation^15^ and the ability of xylem connections to form between parasitic plants and their hosts during the conceptually related process of parasitic plant infection^16^.

The success of wound healing at the graft junction depends on internal factors such as plant hormones and the plant’s developmental stage, but also on external factors such as light intensity^3^, photoperiod^17^ and temperature^18^. In *Arabidopsis*, an elevation in ambient temperature alters various growth and developmental traits such as elongating hypocotyls, petioles and roots^19^. The transcription factor *PHYTOCHROME-INTERACTING FACTOR 4* (*PIF4*) is the major temperature-signaling hub in aerial tissues^20–22^. High temperatures deactivate the photoreceptor Phytochrome B (PhyB) and release its suppression of *PIF4*. The PIF4 protein directly upregulates auxin biosynthesis via activating *YUCCA8* expression^23,24^. In *Arabidopsis*, high temperatures promote a mobile auxin signal^25^ which is activated by epidermal PIF4 in cotyledons^26^. Cotyledon-produced auxin is then transported via the petioles to the hypocotyl where it induces brassinosteroid-induced cell elongation associated with elevated temperatures^25^.

Elevated temperatures during graft healing are used to improve grafting success rates in plants including *Arabidopsis*^3,18^, watermelon^27^, eggplant^4^, walnut^28^, and tomato^29^, however, the molecular basis for the temperature-enhancement of regeneration remain poorly characterised. Here, we investigated the effects of temperature upon various aspects of graft healing including callus formation, tissue attachment and vascular formation and revealed a central role for temperature regulating *PIF4* and *YUCCA2/5/8/9* in leaves to promote vascular formation in grafted stems. Moreover, pharmacological experiments showed that leaf-derived auxin regulated phloem reconnection at the *Arabidopsis* graft junction and xylem bridge formation between the parasite *Phtheirospermum japonicum* (*P. japonicum*) and its host *Arabidopsis thaliana* in a temperature-dependent manner. Taken together, our results suggest a conserved temperature signaling mechanism in leaves regulating vascular regeneration and vascular formation in distant tissues of multiple species.

## Results

### Elevated temperatures enhance vascular formation during grafting

Elevating temperatures during commercial grafting improves success rates^30^ so we tested the effects of temperature upon *in vitro* graft formation in tomato (*Solanum lycopersicum*) and *Arabidopsis*. We applied carboxyfluorescein diacetate (CFDA) to monitor vascular connectivity at the graft junction^8^ (Fig 1A). We observed that grafted tomatoes recovered at 30°C showed significantly faster and higher phloem connection rates compared with those recovered at 20°C (Fig 1B,C). In *Arabidopsis*, we grafted *pSUC2::GFP* scions to wild-type rootstocks (Fig 1A) and observed that the phloem connection rate in *Arabidopsis* was also accelerated by higher recovery temperatures (Fig 1D), similar to tomato grafting. Expression of a reporter gene associated with cambium formation, *DOF6::Venus*^31^, was also enhanced by elevated temperatures (Fig S1A,B). Increasing the recovery temperature from 27°C to 30°C did not promote *Arabidopsis* graft formation, suggesting that 27°C is close to the maximum thermo-induction effect. In contrast, reducing the recovery temperature to 16°C delayed the phloem reconnection rate (Fig 1D). Xylem connectivity assays showed a similar trend since elevating temperatures increased xylem reconnection rates (Fig 1E). We next investigated when and for how long elevated temperatures were required to accelerate graft healing and found that 48 h of warm recovery was sufficient (Fig 1F). However, providing warm temperatures prior to grafting (Fig S1C) had no significant effect upon vascular connectivity (Fig S1D), suggesting that thermo-responsiveness occurs early after wounding and plays an important role during graft healing.

**Fig.1.**
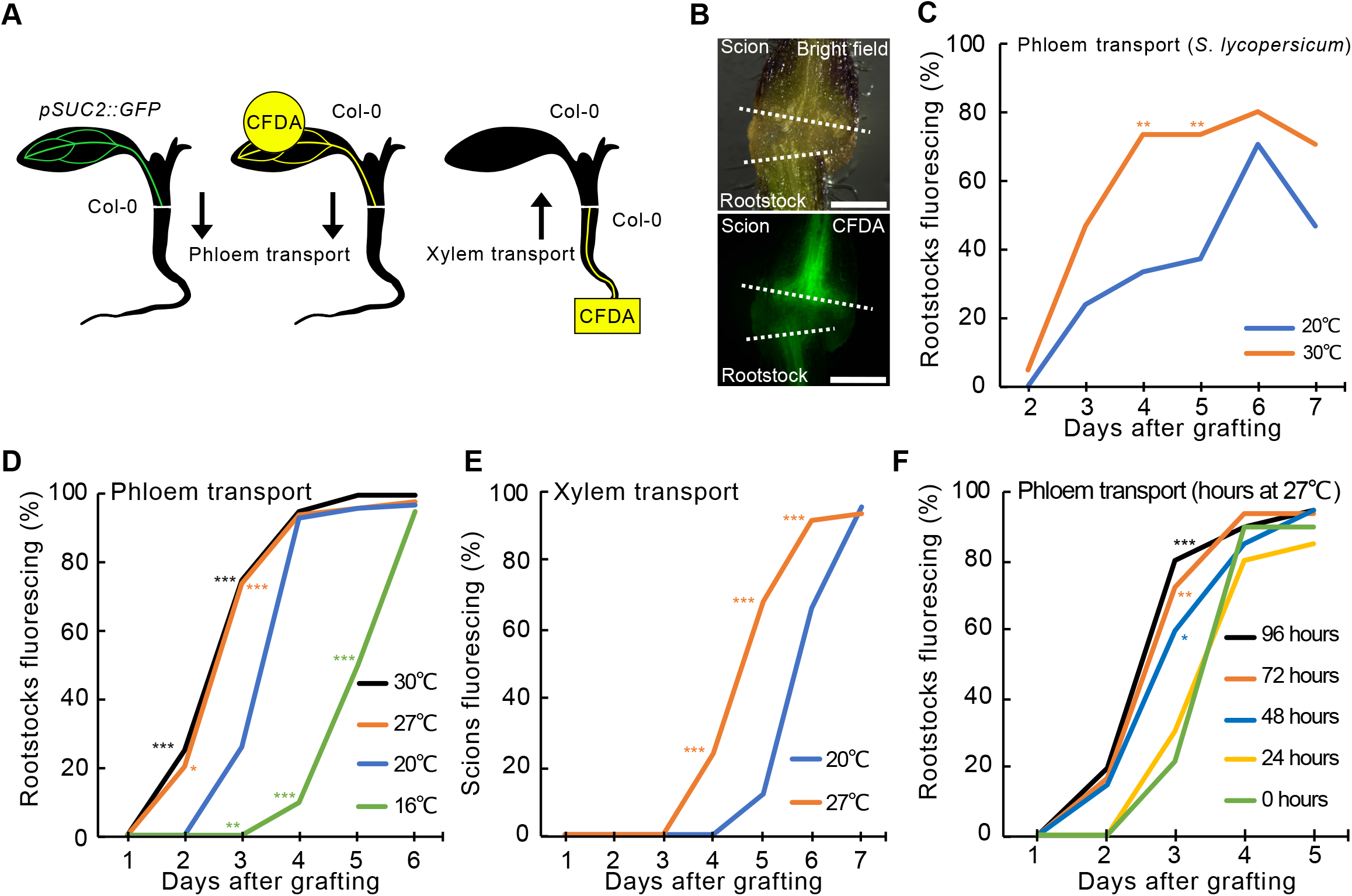
Elevated temperature enhances graft formation. (A) Schematic diagram showing the tissue connection assays used. Phloem transport was measured by detection of fluorescence in the Col-0 rootstock grafted to *SUC2::GFP* scions. Alternatively, fluorescent dye CFDA was applied to scions or rootstocks for the detection of phloem or xylem transport, respectively. (B) Movement of CFDA across the junction of grafted tomato plants. White dashed lines indicate the cut sites at the graft junction. Scale bars = 500μm. (C) Measurement of phloem connection by detection of CFDA in rootstocks of grafted plants recovered at 20°C and 30°C. Values represent proportions of plants with fluorescing rootstocks, n = 24-30 plants per temperature treatment per time point. (D) Phloem reconnection of *Arabidopsis* heterograft *pSUC2::GFP* scions to Col-0 rootstocks recovered at indicated temperatures. Values represent proportions of plants with fluorescing rootstocks, n = 40-80 plants per temperature treatment. (E) Xylem reconnection measurement with CFDA in *Arabidopsis* seedlings grafted and recovered at 20°C and 27°C. Values represent proportions of plants with fluorescing scions, n = 50 plants per temperature treatment and time point. (F) Heterografts of *pSUC2::GFP* scions to Col-0 rootstocks were recovered at 27°C for the indicated period (hour), then transferred to 20°C recovery thereafter. Values represent the proportions of plants with fluorescing rootstocks, n = 40 plants per temperature treatment. (C-F) Individual treatments were compared with the 20°C treatment and asterisks indicate statistical significance (*P<0.05; **P<0.01; ***P<0.001; Fisher’s exact test).

### *PIF4* and *YUCCAs* are required in the cotyledon for temperature-enhanced vascular formation

To better understand how elevated temperatures promoted graft formation, we tested various mutant genotypes associated with temperature response or hormone signaling (Table S1). Most mutants had no effect on temperature enhancement, but the *pif1 pif3 pif4 pif5* quadruple mutant (*pifQ*) and the *pif4* single mutant were exceptional since they did not respond to temperature enhancement at 3 days after grafting (DAG) but had normal grafting dynamics at later time points (Fig 2A; S2A), suggesting they specifically affected temperature enhancement. We tested the spatial requirements of *PIF4* by grafting *pif4* scions to wild-type rootstocks, or vice versa, and observed that *pif4* scions did not respond to the elevated recovery temperature whereas *pif4* rootstocks responded like wild-type (Fig 2A). The cotyledon plays an important role in thermo-sensing^25^ so we generated a graft chimera whereby the cotyledon of *pif4* was initially grafted to a *pSUC2::GFP* shoot and then after graft healing, a hypocotyl graft was performed to a wild-type rootstock for recovery at 20°C and 27°C. Plants with *pif4* cotyledons did not respond to elevated temperatures (Fig 2B), indicating that temperature perception via *PIF4* in the leaves was sufficient for the accelerated graft healing in the hypocotyl. The auxin-biosynthesis gene *YUC8* is a direct target of PIF4^24^ and we found that *YUC8* transcription levels were up-regulated in wild-type plants exposed to elevated temperatures, but down-regulated and non-responsive to elevated temperature in the *pif4* mutant (Fig 2C), consistent with PIF4 acting as an activator of *YUC8*. We also observed that *PIF4* transcript levels were not affected in the *yuc2/5/8/9* quadruple mutant (*yucQ*), supporting the notion that *PIF4* acts upstream of *YUC8*. GUS staining from *pYUC8::GUS* increased in plants grown at 27°C compared to those grown at 20°C and staining was observed mainly in the epidermis, vasculatures, and mesophyll (Fig 2D), consistent with the previously reported expression pattern of *PIF4*^26^. We tested the *yucQ* mutant and found that plants lost grafting thermo-responsiveness when *YUCCA* genes were mutated in the scion (Fig 2E), similar to the *pif4* mutant. The *yucQ* mutant did not affect grafting at later time points indicating its requirement was specific for elevated temperatures. The *yucQ* genotype carries *yuc2, yuc5, yuc8* and *yuc9* mutations yet only *YUC8* was responsive to elevated temperatures (Fig. 2C, S2B), suggesting *YUC8* might play a central role for the observed phenotype. We tested if temperatures would affect the expression of vascular-development genes in the intact seedlings but did not detect any upregulation (Fig S2C), suggestion that wounding and reconnection may be prerequisite for the transcriptional induction. Together, these data indicated a spatial requirement for PIF4 and YUCCA genes for temperature-dependent vascular connectivity.

**Fig. 2.**
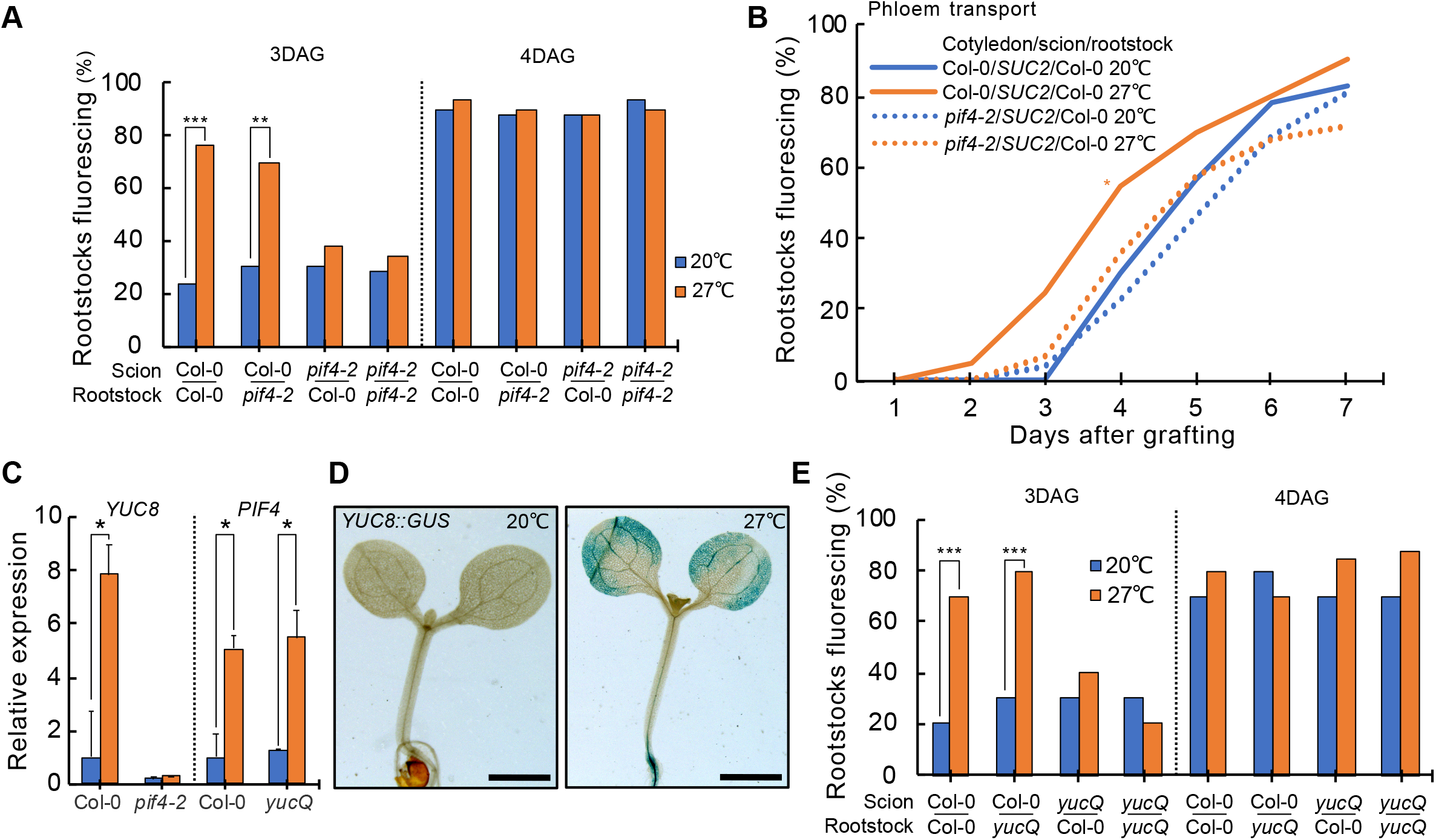
*PIF4* and *YUCCAs* are required in the cotyledon for temperature-enhanced graft formation. (A) CFDA phloem connection rates of plants with *PIF4* mutations in the scion or rootstock recovered at 20°C or 27°C and measured at 3 and 4 days after grafting (DAG). Values represent proportions of plants with fluorescing rootstocks, n = 30-45 plants. (B) *SUC2::GFP* phloem connection rates of three segment cotyledon-hypocotyl grafts with Col-0 or *pif4-2* cotyledons. Values represent proportions of plants with fluorescing rootstocks, n = 40 plants per graft combination and temperature treatment. (C) Relative expression levels of *YUC8* and *PIF4* in Col-0 or *pif4-2* cotyledons incubated at 20°C and 27°C for 48 hours. The values represent the mean(±SD) of three experiments. The asterisk indicates statistical significance (P≤0.01; student’s t-test). (D) GUS histochemical assay of *pYUC8::GUS* transgenic plants grown at 20°C or 27°C for 48 hours. At least 20 seedlings were examined and a representative plant presented. (E) CFDA phloem connection rates of plants with *YUC2/5/8/9* mutations in the scion or rootstock recovered at 20°C or 27°C and measured at 3 and 4 days after grafting (DAG). Values represent the proportions of plants with fluorescing rootstocks, n = 30-45 plants. (A,B,E) Individual treatments were compared with the 20°C treatment and asterisks indicate statistical significance (*P<0.05; **P<0.01; ***P<0.001; Fisher’s exact test).

### Warm temperatures promote vascular formation by enhancing auxin response

Since auxin is important for graft formation and wound healing^7,12,13,32^, we investigated whether auxin contributed to the enhancement of graft formation by elevated temperatures. We applied an auxin transport inhibitor 1-N-naphthylphthalamic acid (NPA) on the petiole to inhibit the transport of auxin from the cotyledon to the hypocotyl, and observed that NPA-treated plants did not respond to temperature enhancement (Fig. 3A), suggesting that cotyledon-derived auxin is essential for this effect. However, graft dynamics of NPA treated plants at 20°C were similar to the controls at 20°C indicating that leaf derived auxin was only relevant for graft formation at elevated temperatures. We next asked whether auxin response at the graft junction was increased by elevated temperatures using the auxin responsive DR5 reporter^33,34^. We found a significant increase of *pDR5::GFP* fluorescence with warm recovery (Fig 3B) indicating that elevated temperatures increased auxin response at the graft junction. Perturbing auxin response in the rootstock with a dominant negative mutant of *BODENLOS/IAA12* (*bdl-2*)^35^ blocked graft formation irrespective of whether grafting was performed at 20°C or 27°C (Fig 3C). However, when *bdl-2* was present in the scion, plants grafted like controls at 20°C but were inhibited in elevated temperature responses at 27°C (Fig 3C). Since we observed earlier that accelerated graft healing in the hypocotyl was due to temperature perception in the leaves, we asked if blocking auxin response in the leaves would also affect the thermo-responsiveness of grafting dynamics. Blocking auxin response in the cotyledon by grafting the *bdl-2* cotyledon to a *pSUC2::GFP* shoot and wild-type rootstock did not affect temperature enhancement (Fig 3D), suggesting that *bdl-2* did not play a role in the leaves for the temperature enhancement of graft formation, and instead, *bdl-2* had its effect at the region of graft junction formation.

**Fig. 3.**
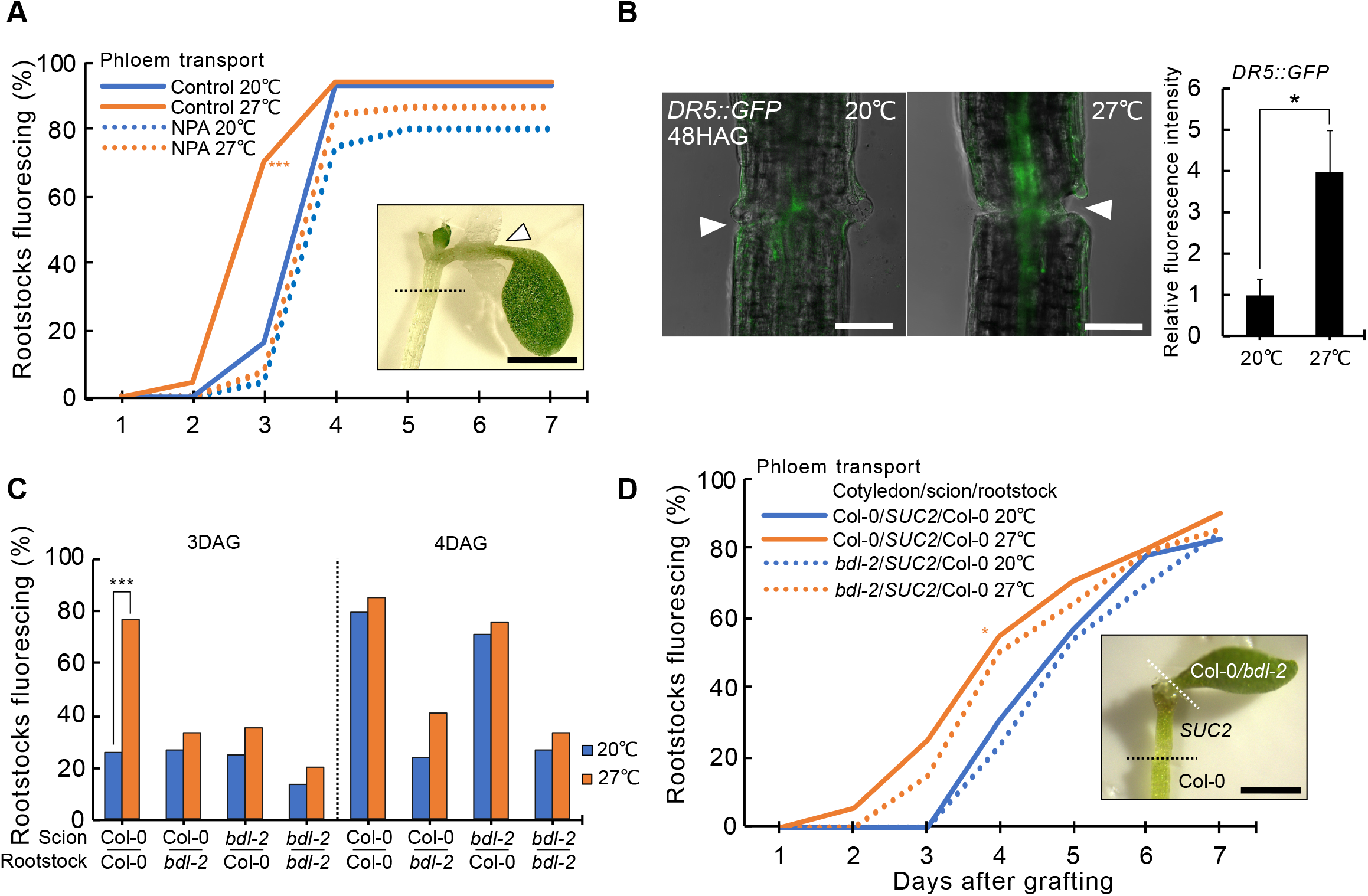
Temperature promotes graft formation by elevating auxin response. (A) *SUC2::GFP* phloem connection rates of plants treated with NPA on the petiole and recovered at 20°C or 27°C. Values represent proportions of plants with fluorescing rootstocks, n = 50 plants per temperature treatment. The white arrow head indicates the site of the NPA plaster. The black dashed line indicates the graft junction of *pSUC2::GFP* scions / Col-0 rootstocks heterograft. Scale bar = 1mm. (B) Detection of fluorescent signals at the graft junction of transgenic plants expressing auxin-responsive *DR5::GFP* 48 hours after grafting (HAG) recovered at 20°C or 27°C. Value indicates the mean from three experiments, each with n = 15-20 plants per temperature treatment. The asterisk indicates statistical significance (P≤0.01; student’s t-test). The white arrow head indicates the graft junction. Scale bars = 100μm. (C) CFDA phloem connection rates of plants with *bdl-2* mutations in the scion or rootstock recovered at 20°C or 27°C and measured at 3 and 4 days after grafting (DAG). Values represent proportions of plants with fluorescing rootstocks, n = 30-45 plants. (D) *SUC2::GFP* phloem connection rates of three segment cotyledon-hypocotyl grafts with Col-0 or *bdl-2* cotyledons. Values represent proportions of plants with fluorescing rootstocks, n = 40 plants. The white and black dashed lines indicate sites of cotyledon and hypocotyl grafting, respectively. Scale bar = 1mm. (A,C,D) Individual treatments were compared with the 20°C treatment and asterisks indicate statistical significance (*P<0.05; **P<0.01; ***P<0.001; Fisher’s exact test).

### Temperature dependent tissue regeneration is widespread

Graft formation involves cell adhesion, callus formation and vascular reconnection^8,36^. Elevated temperatures could affect multiple aspects of this process, so we tested the effects of temperature upon tissue adhesion and callus formation. By picking up plants 1-2 days after grafting with forceps^8^, we could observe that adhesion rates were significantly increased with the elevated temperatures and this enhancement was not affected in *pif4* mutants (Fig S3B). Similarly, using previously described assays^37^ we found that elevated temperatures enhanced callus formation but this enhancement was not affected in *pif4, pifQ, yucQ* or *bdl-2* mutants (Fig 4A, 4B, S3A), suggesting that warm temperatures enhanced multiple aspects of wound healing but that phloem connectivity was specifically dependent on *PIF4* and *YUC2/5/8/9*.

**Fig. 4.**
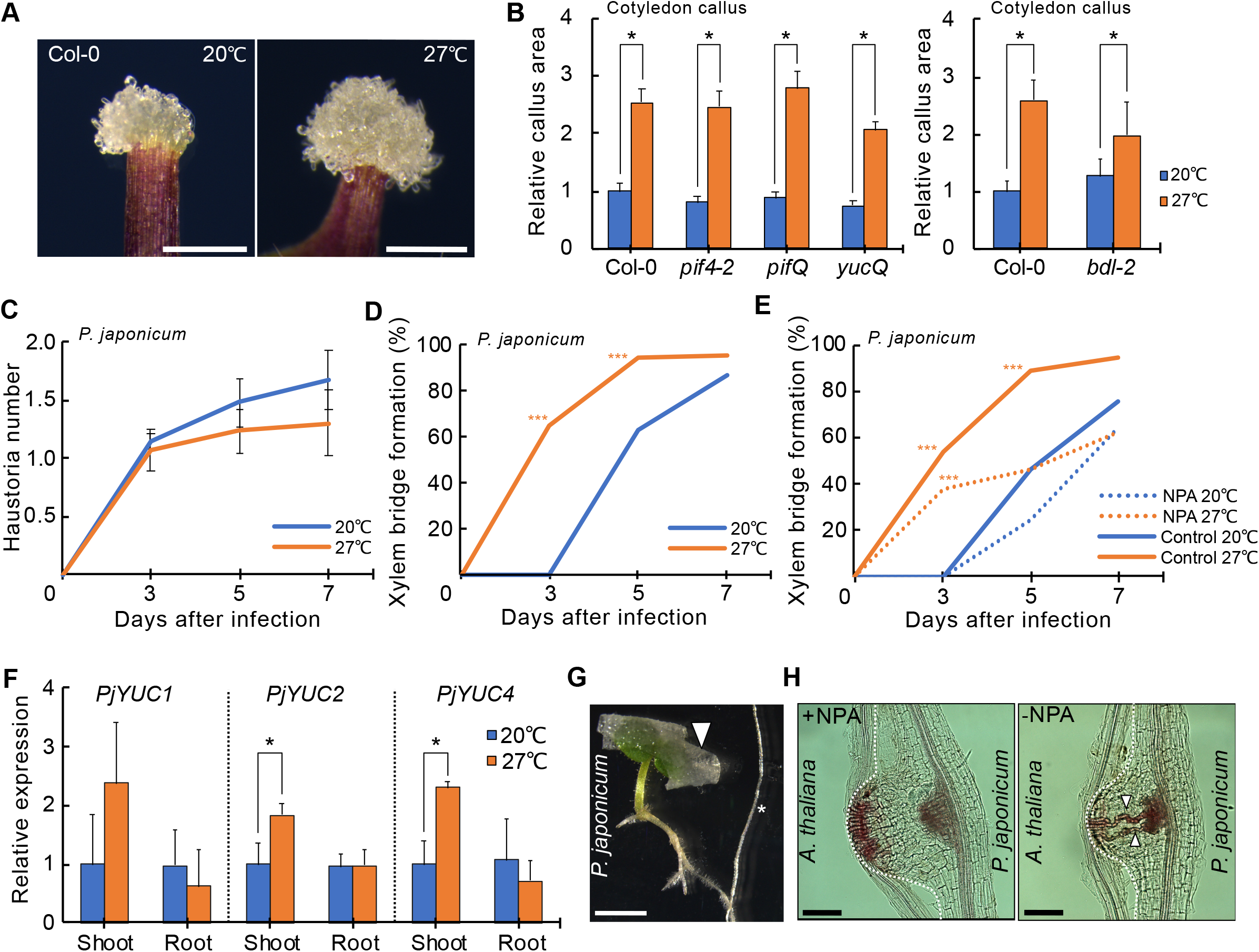
Temperature-dependent tissue regeneration is wide-spread. (A) Callus formation at the wounded site of leaf petioles from 14-day-old Col-0 seedlings and incubated at 20°C or 27°C. Scale bars = 500 μm. (B) Measurement of callus size at 8 days after wounding from various genotypes. Callus size is relative to Col-0 at 20°C and values indicate the mean (±SD) from 60 cotyledons per genotype and temperature treatment. The asterisk indicates statistical significance (P≤0.01; student’s t-test). (C) Haustoria number of *P. japonicum* infecting *Arabidopsis* at 20°C and 27°C. Value indicates the mean(±SD) from four experiments, each with 20 parasitic plants infecting per temperature treatment and time point. (D) Proportion of haustoria forming xylem bridges in *P. japonicum* infecting *Arabidopsis* at 20°C or 27°C. (E) Inhibition of auxin transport from the cotyledons of *P. japonicum* infecting *Arabidopsis* by cotyledon NPA applications at 20°C or 27°C. Values represent proportions of xylem bridge formation, n = 80 (D) and 40 (E) parasitic plant infections per temperature and time point. Individual treatments were compared with the 20°C treatment and asterisks indicate statistical significance (*P<0.05; **P<0.01; ***P<0.001; Fisher’s exact test). (F) Relative expression levels of auxin-related genes in *P. japonicum* at 7 days post infection in shoots and roots during the infection at 20°C or 27°C. Values represent the mean (±SD) of three biological replicates. The asterisk indicates statistical significance (P≤0.01; student’s t-test). (G) Representative images of cotyledon NPA application on *P. japonicum*.The white arrow head indicates the site of the NPA plaster. The asterisk indicates the root of the *Arabidopsis* host. Scale bar = 100 μm. (H) Representative images of haustoria and xylem bridges developed at 27°C with and without NPA applications at 7 DPI. Scale bars = 100μm. The dashed lines outline the interface of *P. japonicum* and *Arabidopsis*. Xylem bridges are indicated by the white arrowheads.

Parasitic plant infections are conceptually similar to grafting^38^ and their infective structures known as haustoria form xylem connections to their hosts to withdraw nutrients. Elevated temperatures are known to increase haustoria numbers^39^ so we tested whether elevated temperatures affected infection of *Arabidopsis thaliana* by the facultative parasite *P. japonicum*. Elevated temperatures during infection did not affect haustoria number (Fig 4C) but we observed that xylem bridge formation was accelerated similar to the effect we observed during xylem reconnection at the graft junction (Fig 1E, Fig 4D). Warm temperatures also increased xylem plate area (Fig S3C) and length (Fig S3D). To investigate the role of leaf-derived auxin, we blocked auxin transport from *P. japonicum* cotyledons using NPA and found that NPA did not affected haustoria number (Fig S3E), but significantly reduced xylem bridge formation at 27°C but did not affect it at 20°C (Fig 4E,G,H). Expression levels of auxin biosynthesis genes *PjYUC2* and *PjYUC4* were up-regulated by elevated temperatures in *P. japonicum* shoots but not roots (Fig 4F). Thus, similar to grafting, shoot-derived auxin contributed to vascular formation in basal tissues of *P. japonicum*.

## Discussion

Elevated temperatures have dramatic effects upon both animal and plant development^40–42^ and here we demonstrate that warm temperatures enhanced multiple aspects of wound healing including tissue adhesion, callus formation and vascular regeneration that we could mechanistically separate based on their dependency on *PIF4*. *PIF4* was specifically required in developing leaves to promote vascular regeneration, and this protein is known to activate auxin biosynthesis^23^, suggesting that cotyledon-derived auxin was sufficient to enhance vascular formation at the graft junction (Fig S4). Notably, cotyledon-derived auxin did not appear necessary for successful graft formation at ambient temperatures consistent with previous findings that polar auxin transport did not enhance auxin response at a wound site^14^ and removing cotyledons in *Arabidopsis* and tomato grafts did not impair grafting success^17,43,44^. One interpretation of our results is that cotyledon-derived auxin can modify the rate of regeneration but not the final outcome, and instead, cotyledon photosynthetic capacity might be more important since grafting in the absence of any cotyledons requires exogenous sugars for success^17^. Enhancing auxin response at the graft junction likely enhanced graft healing through auxin’s known roles in promoting xylem differentiation and activating cambium likely in part by including *DOF6* (Fig S1)^45,46,47^, processes that were likely affected in the auxin resistant *bdl-2* scion^35^. However, *bdl-2* rootstocks did not respond to temperature and inhibited grafting regardless of temperature indicating that the rootstock had a different requirement for auxin response and appeared more sensitive to auxin perturbations. Callus formation and tissue attachment were *PIF4* independent and it’s possible that other hormones play a role in these processes such as cytokinin which is critical for hypocotyl callus formation^9^ and brassinosteroids that are involved in temperature-dependent callus formation^2^.

Enhanced temperatures also accelerated haustoria development in *P. japonicum*. This effect was specific to xylem bridge formation and we did not observe differences in haustoria numbers at elevated temperatures. Previous reports demonstrated that blocking auxin production prevents haustoria formation whereas inhibiting auxin transport prevents xylem bridge formation^16^. We extended a role for auxin and found that cotyledon-derived auxin played an important role in responding to elevated temperatures to enhance xylem bridge formation. Our observations in grafted plants and parasitic plants demonstrates a common mechanism by which temperature sensing in leaves changes vascular development and contributes to modifying the rate of vascular regeneration or vascular formation. Such modulations could provide developmental plasticity in response to environmental changes and confer a fitness advantages of earlier regeneration or earlier differentiation of vascular tissues to promote water and photosynthate transport. High temperatures are also known to enhance regeneration of *Hydra* tentacles^48^, zebrafish fin^49^, and flatworm testes^50^ suggesting the enhancement of regeneration by elevated temperatures is universally relevant and an important aspect to modify to improve regeneration.

## Material and methods

### Plant materials, growth conditions, and grafting

*Arabidopsis thaliana* (L.) ecotypes Columbia (Col-0) was used throughout this study unless otherwise indicated. Mutant lines used included *pif4-2* (CS66043), *pifQ* (CS66049), *yucQ* (CS69869), *bdl-2*^35^.The previously published transgenic lines include *pSUC2::GFP*^51^, *pYUC8::GUS*^52^, *pDR5rev::GFPer*^33^, and *pDOF6::VENUSer*^31^. For *in vitro* germination, seeds were surface sterilized with 70% (v/v) ethanol for 10 minutes, followed by 90% (v/v) ethanol for 10 minutes. The seeds were then sown and germinated on ½MS media (1% plant agar), pH 5.8. After stratification in the dark at 4°C overnight, the seeds were transferred to 20°C short-day growth conditions (8 h of 140 μmol m^−2^ s^−1^). *Arabidopsis* grafting was carried out according to previously published protocols^43,53^, and recovered at 16°C, 20°C, 27°C or 30°C. For the three-segment cotyledon-hypocotyl grafting, cotyledon grafting was first performed when plants were four days old^3^, then after three days of recovery at 20°C, the attached plants were used for the hypocotyl grafting.

MoneyMaker tomato (*S. lycopersicum*) seeds were sterilized in 75%bleach solution for 20 minutes, then rinsed at least five times with sterile water. The seeds were then sown on ½ MS media (1% agar) and germinated at 25°C under short-day conditions (8 h of 140 μmol m^−2^ s^−1^). Tomato grafting was performed using seven-day-old seedlings. A straight cut was made in the middle of the hypocotyl using a scalpel. Rootstocks and scions were held together within a silicone tube (0.8 mm diameter). The grafted seedlings were transferred on 1% agar media and grown under short-day conditions (8 h of 140 μmol m^−2^ s^−1^), at either 20°C or 30°C.

### Phloem and xylem connection assays

Xylem and phloem connections were monitored by the movement of the fluorescent dye carboxyfluorescein diacetate (CFDA) (Thermo Scientific) across the graft junction. To measure phloem connection, the cotyledon of the *Arabidopsis* grafted plants was wounded with forceps, and then CFDA solution (1mM) was applied on the surface using a pipette. After 1 h incubation at room temperature, fluorescent signals in the rootstocks were detected. Alternatively, *pSUC2::GFP*^51^ scions were grafted to wild-type rootstocks, and the GFP signals in the roots were observed daily. For the xylem connection assay, a previously published protocol was modified slightly^3^. In brief, grafted plants with cut root tips were place on a piece of Parafilm, then 1 μl of 1 mM CFDA solution was dropped on the cut site. The signals in the cotyledons were detected after 1 h. For tomato phloem assays, one of the two cotyledons was cut and a drop of CFDA (5 mM CFDA in 1% agar) was applied on the cut site. Seedlings were kept in the dark for at least 2 h. Transversal sections of the hypocotyl (at the shoot-root junction) were made at 2 h and placed on slides to help monitor fluorescence movement. *Arabidopsis* plants and tomato sections were observed under a Leica M205 FA microscope, with GFP filter to detect CFDA fluorescence in the phloem or xylem.

### Parasitic plant infection assays

*Phtheirospermum japonicum* (*P. japonicum*) seeds were surface sterilised by washing with 70% ethanol for 20 minutes, followed by 95% ethanol for 5 minutes, and sown on ½ MS with 1% sucrose and 0.8% agar. After stratification at 4°C in darkness overnight, the plates were moved to a growth cabinet at 20°C in short day conditions (8 hr of 140 μmol m^−2^ s^−1^). Four-day-old *P. japonicum* seedlings were transferred to 0.8% water agar for starvation prior to infection. For the infection, the root of a five-day-old Arabidopsis seedlings was aligned to each *P. japonicum* root, and the infection setup was incubated at 20°C or 27°C in short day conditions (8 h of 140 μmol m^−2^ s^−1^). At 24 h post infection, swellings on *P. japonicum* root corresponding to early haustoria were marked as day-1 haustoria. The number of haustoria and presence of a xylem bridge were quantified using a Zeiss Axioscope A1 microscope at 3, 5- and 7-days post infection. Xylem plate area and length were measured on 7 DPI haustoria stained with Safranin-O using a previously published protocol^54^.

### Histological staining and confocal imaging

Histological staining of GUS was analyzed in the 8-day-old transgenic *pYUC8::GUS* seedlings. Seedlings were incubated for 12 h at 37°C with the substrate solution (1 mM 5-bromo-4-chloro-3-indolyl-β-D-glucuronide, pH 7.0, 100 mM sodium phosphate buffer, 10 mM Na2EDTA, 0.5 mM potassium ferricyanide, 0.5 mM potassium ferrocyanide, and 0.1% Triton X-100). Stained seedlings were washed with 70% ethanol overnight to remove chlorophyll, and were then photographed with a Leica M205 FA microscope. For confocal microscopy, fluorescent images of the graft junctions of transgenic seedlings expressing fluorescent proteins were taken on Zeiss LSM-780 laser scanning confocal microscope. The images were processed and analyzed using FIJI software (version 2.1.0/1.53c).

### NPA treatment assay

The application of the auxin inhibitor 1-N-naphthylphthalamic acid (NPA) plasters on *Arabidopsis* was adapted from a previous study^25^. Plants were grown in short-day conditions at 30 μmol m^−2^ s^−1^ to induce longer petioles prior grafting for a more efficient application of NPA plasters. Thin strips of cellulose tissue were soaked in a lukewarm agar solution (1 %) with or without 100 μM NPA (Duchefa) and were carefully positioned across petioles using fine forceps after grafting. For the application of NPA on *P. japonicum*, the NPA plasters were placed on the cotyledons just before the infection assay.

### Plant attachment and callus formation assays

For the attachment assays, grafted *Arabidopsis* recovered at 20°C to 27°C were picked up with forceps at the root/hypocotyl junction and scions scored whether they remained attached or fell apart. The petiole callus formation^37^ assays was adapted from previously published protocols. The explants were incubated at 20°C to 27°C and the area of callus was quantified by ImageJ (version 2.1.0/1.53c). Callus induction was quantified as a percentage of explants with more than one callus cell developing from wound sites.

### Gene expression analyses

For *Arabidopsis*, total RNA was extracted from whole seedlings or cotyledons using ROTI Prep RNA MINI (Roth), then subsequently treated with DNase I (NEB) to eliminate DNA contamination. The cDNA was synthesized with Maxima First Strand cDNA Synthesis Kit for RT-qPCR (Thermo Scientific). The transcript levels were measured by quantitative real-time PCR (qPCR) using SYBR-Green master mix (Applied 512 Biosystems) with specific primers (Table S2). The data were normalized against *PP2A*^55^. The relative expression was calculated using Pfaffl method^56^. All reactions were carried out with three biological replicates, each with three technical replicates. For *P. japonicum*, seven-day-post-infection plants were separated from host *Arabidopsis*, and the shoot and root samples were collected. Total RNA extraction, cDNA synthesis, and qPCR were performed using the mentioned protocol with *P. japonicum* specific primers (Table S2). The data were normalized against *PjPP2A*^57^.

### Statistics

For pairwise comparisons of frequencies, Fisher’s exact test was used with the indicated sample sizes. For pairwise comparisons of continuous data, student’s t-test was performed.

## Supplemental data

Supplemental figure S1

Supplemental figure S2

Supplemental figure S3

Supplemental figure S4

Supplemental table S1: Genotypes tested for phloem connection. 7-day-old seedlings were self-grafted (unless indicated) and recovered at 20°C and 27°C. Phloem connection measurement was done by CFDA assay at 3 and 4 DAG. The value indicates percentage of rootstocks fluorescing of at least 20 plants. Asterisks indicate the heterograft with Col-0 rootstocks.

Supplemental table S2: Primer sequences used for qPCR

## Acknowledgements

We thank Phil Wigge for help initiating the project. We thank the Nottingham Arabidopsis Seed Centre for providing seeds. We thank Valentin Codemard for contributing to drawing the temperature perception model. PTS and CWM were supported by a Wallenberg Academy Fellowship (2016-0274). ML, CM and CWM were supported by an ERC starting grant (GRASP-805094). MQ is supported by the Deutsche Forschungsgemeinschaft (Qu 141/3-2).

