## Supplemental Figures 1-4 for "High temperature perception in leaves promotes vascular regeneration in distant tissues"

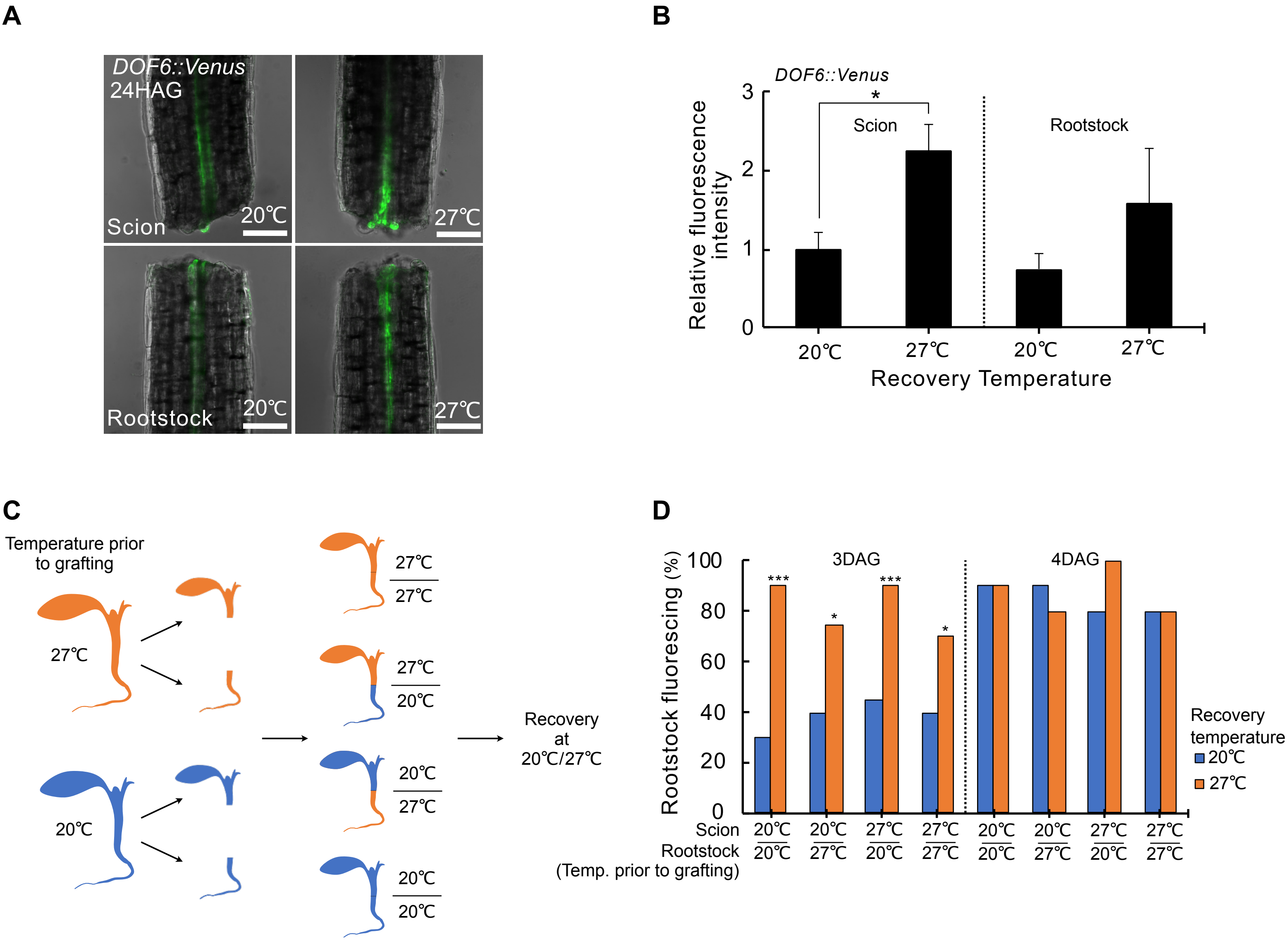


**Supplemental figure S1:** (A, B) Confocal images of *DOF6::Venus* scions and rootstocks of plants recovered at 20℃ or 27℃ at 24 hours after grafting (HAG). Plants were grafted but not firmly attached during imaging at 24 hours so scions and rootstocks are shown separately. The values represent the mean(±SD) of two experiments, each with 15-20 plants per temperature treatment. The asterisk indicates statistical significance ($P \leq 0.01$; student's t test). Scale bars= 100 μm. (C) Col-0 seedlings cultivated for 7 days at 20℃ or 27℃ were cut and grafted with various temperature combinations. The grafted plants were subsequently transferred to recover at 20℃ or 27℃. (D) Measurement of phloem connection by CFDA assays in grafted Col-0 plants recovered at 20℃ and 27℃, at 3 and 4 days after grafting (DAG). The combination of temperature treatment prior to grafting is indicated. The values represent the proportions of plants with fluorescing rootstocks, n = 20-30 plants. The individual treatments are pairwise compared with the 20℃/20℃ graft combination at 20℃. Asterisks indicate statistical significance (*$P$<0.05; **$P$<0.01; ***$P$<0.001; Fisher's exact test).

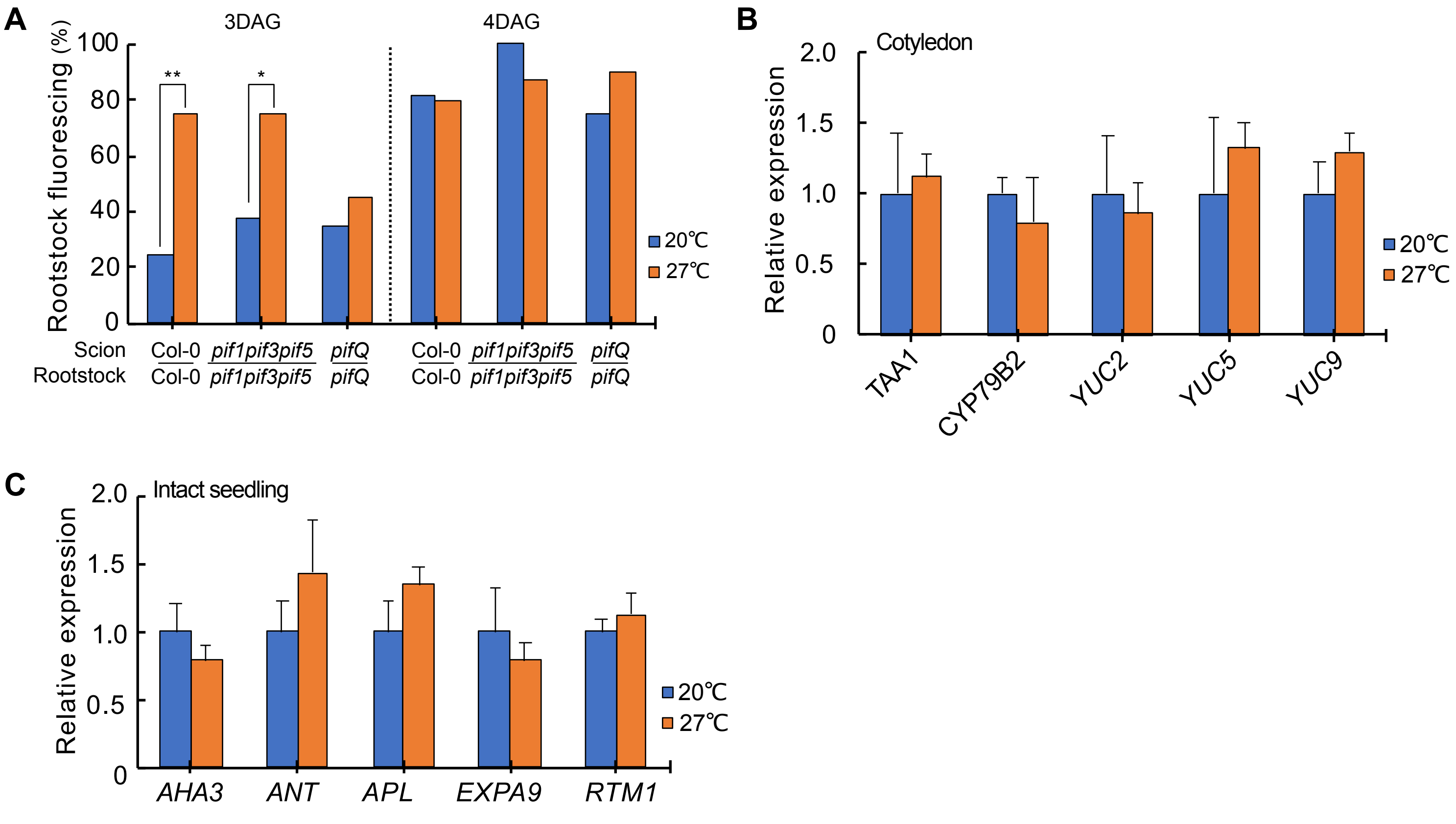


**Supplemental Fig S2** CFDA assay phloem connection rates of plants with *pif1pif3pif5* or *pif1pif3pif4pif5* (*pifQ*) mutations recovered at 20℃ or 27℃, at 3 and 4 days after grafting (DAG). The values represent the proportions of plants with fluorescing rootstocks, n=20. The asterisks indicate statistical significance (**P*<0.05; ***P*<0.01; ****P*<0.001; Fisher's exact test). (B) Relative expression levels of selected auxin biosynthesis genes in the cotyledons of 7 day-old Col-0 incubated at 20℃ and 27℃ for 48 hours. (C) Relative expression levels of selected vascular genes in 7 day-old Col-0 seedlings incubated at 20℃ and 27℃ for 48 hours. (B,C) The expression levels of each gene is normalized to expression at 20℃. The values represent the mean(±SD) of three biological replicates. Student's t-test was performed (*P*≤ 0.01), and the data show no significant difference compared to 20℃.

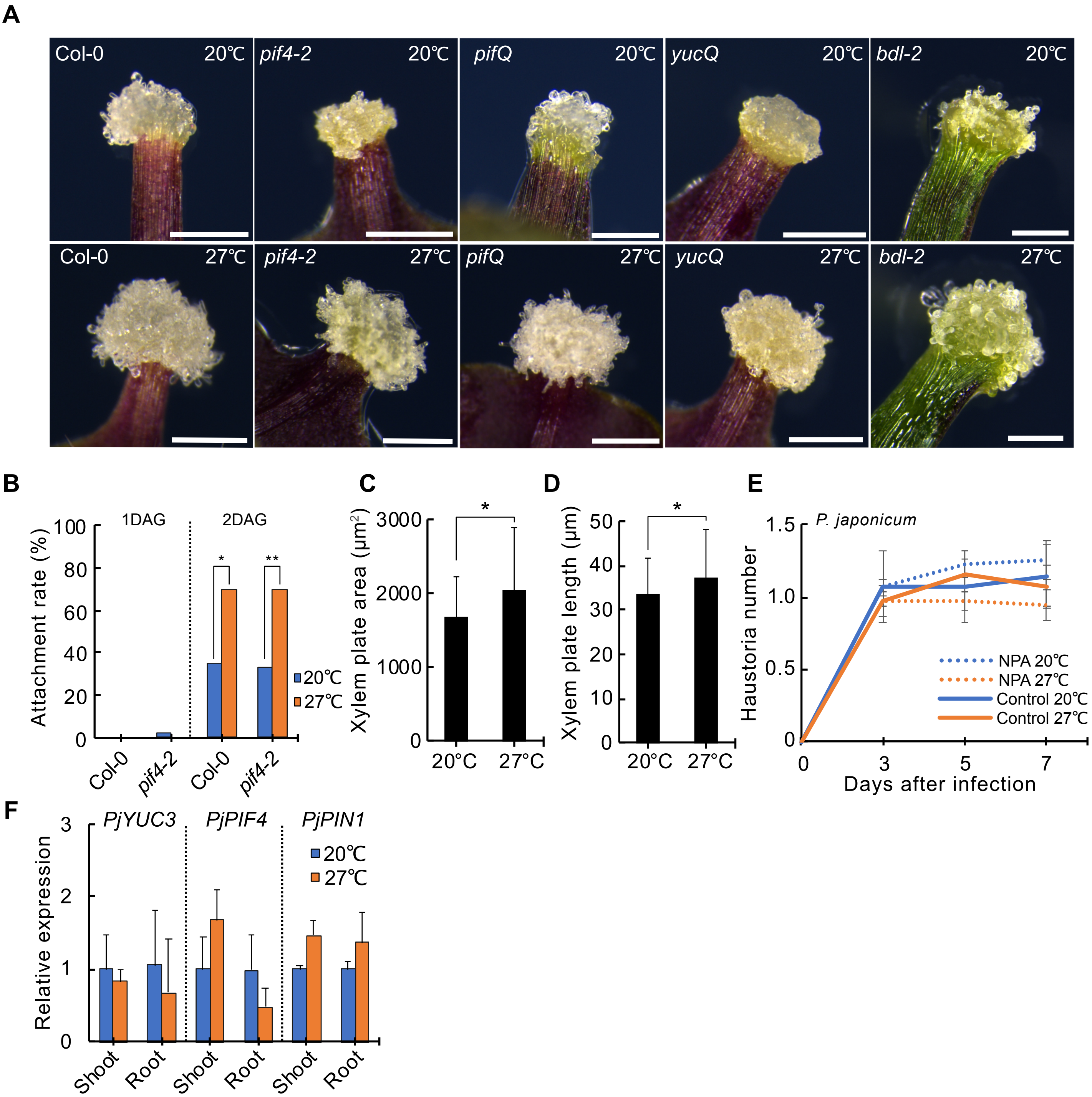


**Supplemental Fig S3** Callus formation at the wounded site of petioles at 8 days after wounding. Leaves were taken from 14 day-old seedlings and incubated at 20℃ or 27℃. Scale bars= 500 µm. The images are representative of the average of 60 leaves from each genotype and temperature treatment. Col-0 taken from Fig.4A. (B) Plant attachment rate in grafted plants recovered at 20℃ and 27℃ at 1-2 days after grafting (DAG). The values represent the proportions of attached plants, n=30-45. The asterisks indicate statistical significance (**P*<0.05; ***P*<0.01; ****P*<0.001; Fisher's exact test). (C-D) Measurement of xylem plate area (C) and length (D) in the haustoria of the parasitic plant *P. japonicum* infecting the host plant *Arabidopsis* at 7 days post infection (DPI) at 20℃ or 27℃. The value indicates the mean (±SD) of 74-75 haustoria per temperature treatment. The asterisk indicates statistical significance (p≤0.01; Student's t-test). (E) Inhibition of auxin transport from the cotyledon of the parasitic plant infecting the host by cotyledon NPA applications at 20℃ and 27℃. The value indicates the mean(±SD) from four biological replicates, each with 20 parasitic plants infection per temperature treatment per time point. Student's t-test was performed (*P*≤ 0.01), and the data show no significant difference against 20℃. (F) Relative expression levels of auxin-related genes in the parasitic plants at 7 DPI in the shoots and roots during the infection at 20℃ or 27℃. The values represent the mean(±SD) of three experiments. The asterisk indicates statistical significance (p≤0.01; Student's t-test).

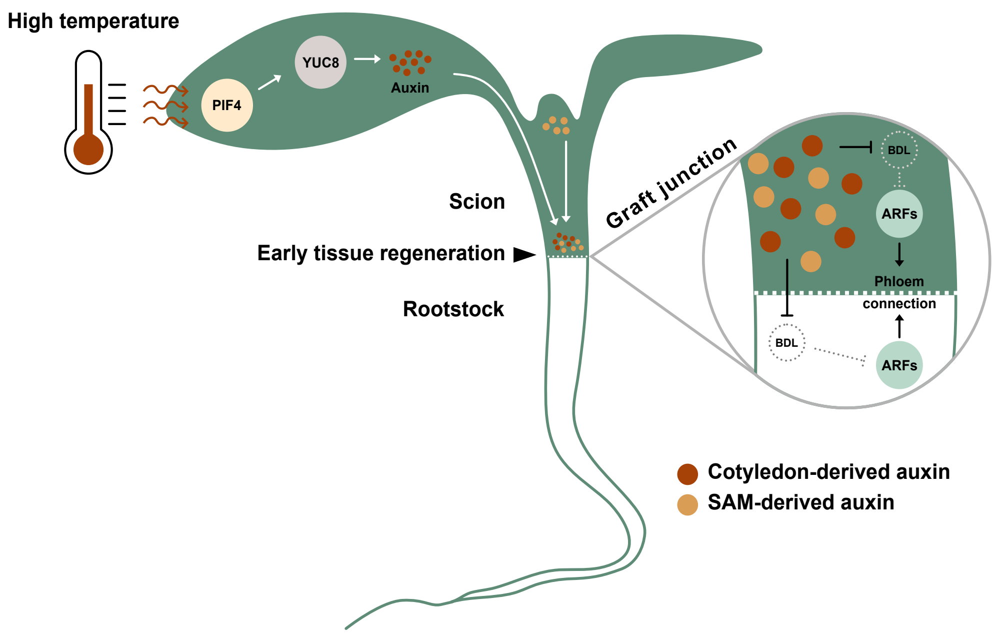


**Supplemental Fig S4** A proposed model for temperature-dependent vascular regeneration. Elevated temperatures cause PIF4 accumulation in the cotyledons and activates *YUC8* expression to catalyze auxin production. The cotyledon-derived auxin is subsequently transported through the petiole to the hypocotyl graft junction where it degrades the repressor BDL (IAA12). ARFs are released to activate auxin responses that lead to phloem reconnection. Auxin also moves across the graft junction to activate the auxin response from the rootstocks[9]. Shoot apical meristem derived auxin likely contributes to graft formation regardless of temperature.
